# β-Arrestin2 Drives Sex-Specific Akt/GSK3β Signaling, Aβ Pathology, and Cognitive Decline in Alzheimer’s Disease Mice

**DOI:** 10.64898/2026.08.10.743981

**Authors:** Fatemeh A. Panahi, Fatemeh Babaei, Tash-Lynn L. Colson, Stephen S. G. Ferguson

**Affiliations:** University of Ottawa Brain and Mind Institute; Departments of Cellular and Molecular Medicine, University of Ottawa, Ottawa, Ontario, K1H 8M5, Canada

## Abstract

Sex is a major determinant of Alzheimer’s disease risk and progression, yet the molecular mechanisms underlying this dimorphism remain poorly defined, limiting the development of sex-informed therapeutics. β-Arrestin2 is a pervasive, multifunctional regulator common to a host of G protein-coupled receptors (GPCRs) in the brain, but whether it has a sex-dependent role in Alzheimer’s disease is unknown. Here, we demonstrate that β-arrestin2 deficiency produces sexually dimorphic effects on Aβ pathology, neuroinflammation, cognition and autophagic flux in APPswe/PS1ΔE9 (APP/PS1) mice. In males, *Arrb2* deletion reduced Aβ oligomer burden, enhanced autophagy, suppressed astrocytic and microglial reactivity, and broadly rescued cognition encompassing spatial working memory, spatial learning, cognitive flexibility, and recognition memory. In females, Aβ pathology and astrogliosis was unchanged and microgliosis was enhanced, with cognitive improvement limited to recognition memory. The male-specific reduction in pathology was accompanied by decreased S473-Akt and S9-GSK3β phosphorylation and enhanced GSK3β/ZBTB16-mediated autophagy, identifying β-arrestin2 as a molecular switch driving sex-restricted Aβ pathology, glial activation, and cognitive decline in male APP/PS1 mice. These findings identify β- arrestin2 as a sex-dependent node linking Aβ pathology to cognitive outcomes in males but not females, underscoring the necessity of sex-stratified consideration in the design of GPCR-targeted Alzheimer’s disease therapeutics.

## Introduction

Alzheimer’s disease (AD) is a neurodegenerative disorder marked by severe memory loss and cognitive decline that predominantly affects persons over the age of 65^1–3^. There are currently no truly effective treatments to slow or cure AD progression, and with an aging global population, AD has the potential to become an epidemic healthcare burden. AD also disproportionately affects women, yet sex has been frequently neglected as a biological variable in preclinical research, and the molecular basis for this sex difference remains poorly defined; a gap that limits the development of sex-informed therapeutics^4–7^.

Several G protein-coupled receptors (GPCRs) have been implicated in AD pathology, including the angiotensin type 1A receptor (AT1AR)^8^, G protein receptor 3 (GPR3)^9,10^, corticotropin-releasing factor receptor 1 (CRFR1) ^11^, and β1 adrenergic receptor (β1AR)^12^. Despite their structural diversity, these receptors converge on a common downstream mechanism, with β- arrestin serving as the shared regulatory protein linking receptor activation to Aβ pathology^13^. This convergence supports the hypothesis that β-arrestins, rather than any single receptor, function as a central regulatory node linking GPCR activity to AD pathogenesis.

β-Arrestins serve as central regulators of GPCR function throughout the brain, acting broadly across receptor subtypes to shape the signaling output of neuronal GPCRs^14–16^. First identified as cofactors for G protein-coupled receptor kinase 2 (GRK2)-mediated desensitization of the β2-adrenergic receptor^17^, β-arrestins also serve as adaptors for GPCR endocytosis and as signaling scaffolds capable of sustaining downstream signaling independent of continued receptor activation, giving rise to the concept of biased agonism through G protein- versus β-arrestin-dependent signaling^18–24^. Both β-arrestin1 and 2 isoforms are highly expressed in brain, though β- arrestin1 predominates over β-arrestin2 by up to 10-fold; only β-arrestin2 localizes to the postsynaptic density, consistent with a distinct role for this isoform in synaptic and disease-relevant signaling^25,26^.

β-Arrestin2 has been directly implicated in neurodegenerative disease pathogenesis in AD, Parkinson’s disease, and frontotemporal dementia^27–30^. β-Arrestin2 is elevated in both FTD and AD patient brain, while β-arrestin1 mRNA is reduced, and *Arrb2* variants associate with late-onset AD risk^27,28,30,31^. Functionally, *Arrb2* deletion reduces tau pathology and rescues synaptic deficits in P301S mice and separately reduces Aβ42 production through disruption of a GRK-dependent interaction with the γ-secretase subunit APH1A^28,32^. Astrocyte-specific *Arrb2* knockdown similarly ameliorates Aβ-induced cognitive deficits^10^. However, despite this established, causal role across multiple AD-relevant pathologies, whether β-arrestin2 also contributes to AD’s pronounced sex bias has not been examined.

This is a notable gap, given existing evidence for sex-specific GPCR signaling in AD models. An mGluR5 negative allosteric modulator rescues cognition and reduces Aβ pathology selectively in male APP/PS1 mice, acting via GSK3β/ZBTB16-mediated autophagy triggered by dephosphorylation of GSK3β at Ser9^33–35^, and CRFR1 overexpression produces AD-like pathology with its own sex-specific pattern^11,36^. β-Arrestin2 is required for mGluR5-dependent synaptic plasticity and for Akt activation, including downstream of D2 dopamine receptor signaling^37–40^. In Huntington disease mice Akt drives inhibitory Ser9 phosphorylation of GSK3β and suppresses GSK3β/ZBTB16-dependent autophagy^41–43^ and β-arrestin2 separately promotes Aβ42 production via γ-secretase regulation in AD mice^27,30,32^. Whether β-arrestin2 links these events to regulate autophagy, and whether either mechanism operates in a sex-specific manner, has not been tested. Here, we show that Arrb2 deletion reduces Aβ plaque burden and soluble Aβ selectively in male APP/PS1 mice, with corresponding reductions in astrogliosis and microgliosis and rescue of spatial working memory, learning, and cognitive flexibility deficits predominantly in male mice. This male-specific reduction in pathology is accompanied by decreased pS473-Akt and pS9-GSK3β and enhanced GSK3β/ZBTB16-mediated autophagy, identifying β-arrestin2 as a molecular switch driving sex-restricted Aβ pathology, glial activation, and cognitive decline in APP/PS1 mice.

## Results

### *Arrb2* deletion results in sex-specific reduction of Aβ pathology i APP/PS1 mice

Amyloid-β plaque deposition is a defining pathological hallmark of APP/PS1 mice, and prior work implicates β-arrestin2 in Aβ production and clearance^10,27,30,32^. We therefore asked whether *Arrb2* deletion-dependent alterations in Aβ pathology differ by sex. At 9 months of age, both male and female APP/PS1 mice exhibited extensive β-amyloid plaque deposition in both the cortex and hippocampus (Fig. 1A and B). *Arr2b* deletion resulted in a significant decrease in β- amyloid plaque density in the cortex of male but not female APP/PS1 mice (Fig. 1A-D). However, at this age, *Arr2b* deletion was not found to alter β-amyloid plaque density in either the dentate gyrus or CA1 regions of the hippocampus (Fig. 1A-D), although there was a trend toward increased β-amyloid plaque density in the dentate gyrus of female animals lacking β-arrestin2 expression (Fig. 2D). Soluble Aβ42 aggregates were significantly increased in cortical tissue from 9-month-old male and female APP/PS1 mice when compared to WT mice and *Arr2b* deletion in male but not female APP/PS1 mice reduced soluble Aβ42 aggregates to WT levels (Fig. 1E). Thus, *Arr2b* deletion results in male-specific reductions in Aβ pathology in APP/PS1 mice.

**Fig. 1:**
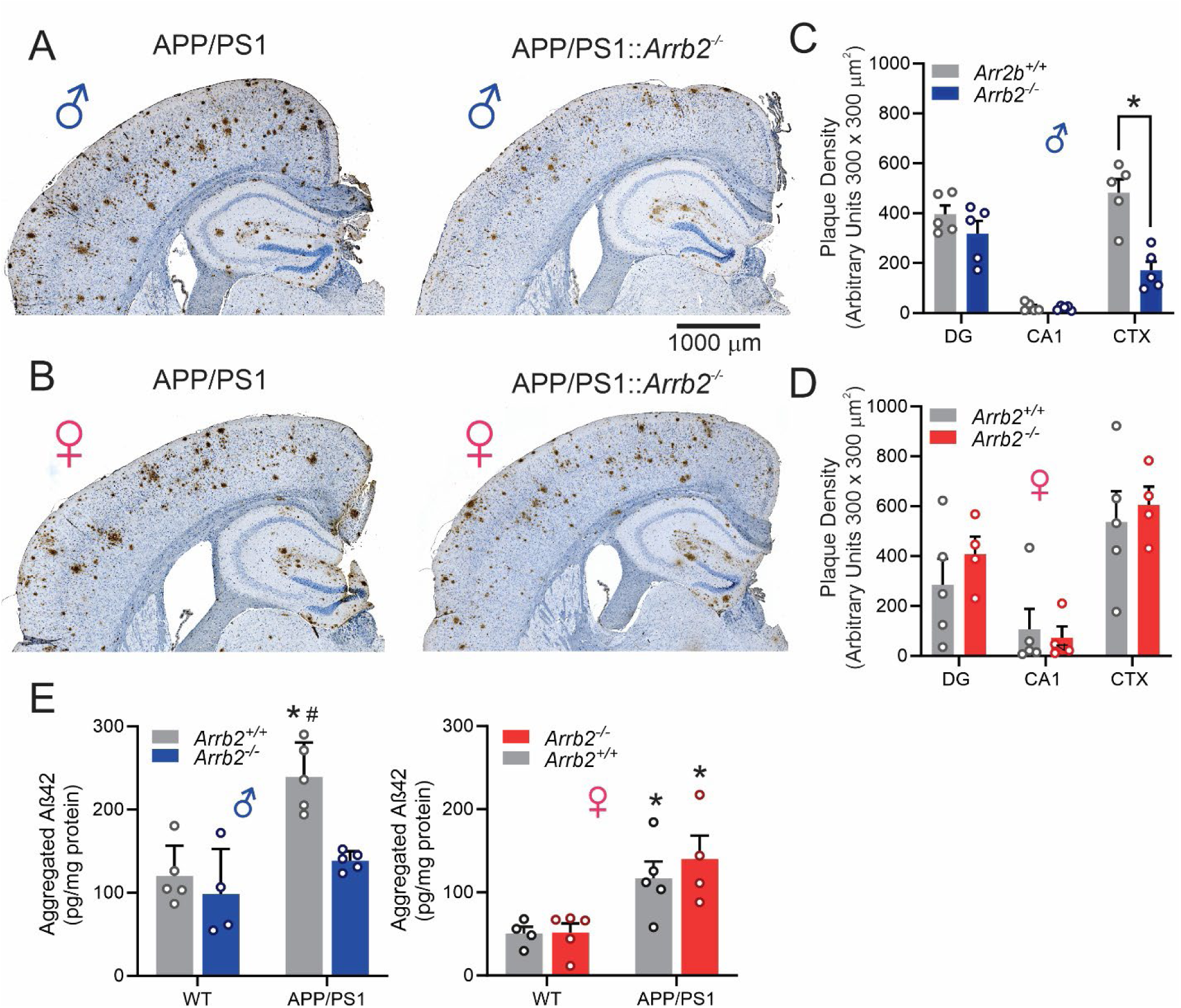
Effect of *Arrb2* deletion on Aβ pathology in male and female wild-type and APP/PS1 mice. **(A)** Representative images of Aβ staining of prefrontal cortex and hippocampus from male APP/PS1 and APP/PS1;*Arrb2^-/-^* mice. **(B)** Representative images of Aβ staining of prefrontal cortex and hippocampus from female APP/PS1 and APP/PS1;*Arrb2^-/-^* mice. **(C)** Quantification of plaque density in 2-4 300 x 300 µm regions of the dentate gyrus (DG), CA1 and cortex (CTX) of six brain slices per male APP/PS1 (*Arrb2^+/+^*) and APP/PS1;*Arrb2^-/-^* (*Arrb2^-/-^*) mouse. Data are presented as mean ± SEM (n =5 mice per group). Statistical analysis was assessed by two tailed paired T-test. *<0.05 versus corresponding region in *Arrb2^+/+^* mice. **(D)** Quantification of plaque density in 2-4 300 x 300 µm regions of the dentate gyrus (DG), CA1 and cortex (CTX) of six brain slices per female APP/PS1 (*Arrb2^+/+^*) and APP/PS1;*Arrb2^-/-^* (*Arrb2^-/-^*) mouse. Data are presented as mean ± SEM (n = 4-5 mice per group).Statistical analysis was assessed by two tailed unpaired T-test. *<0.05 versus corresponding region in *Arrb2^+/+^*mice. **(E)** Aβ oligomer concentrations (pg/mg) from male and female WT, *Arrb2^-/-^*, APP/PS1, and APP/PS1;*Arrb2^-/-^* mouse cortices. Data are presented as mean ± SEM (n = 4-5 mice per group). Statistical analysis was assessed using one-way ANOVA for male mice and for female mice individually followed by Tukey’s multiple comparisons test. *p<0.01 versus WT (Arrb2^+/+^).

**Fig. 2:**
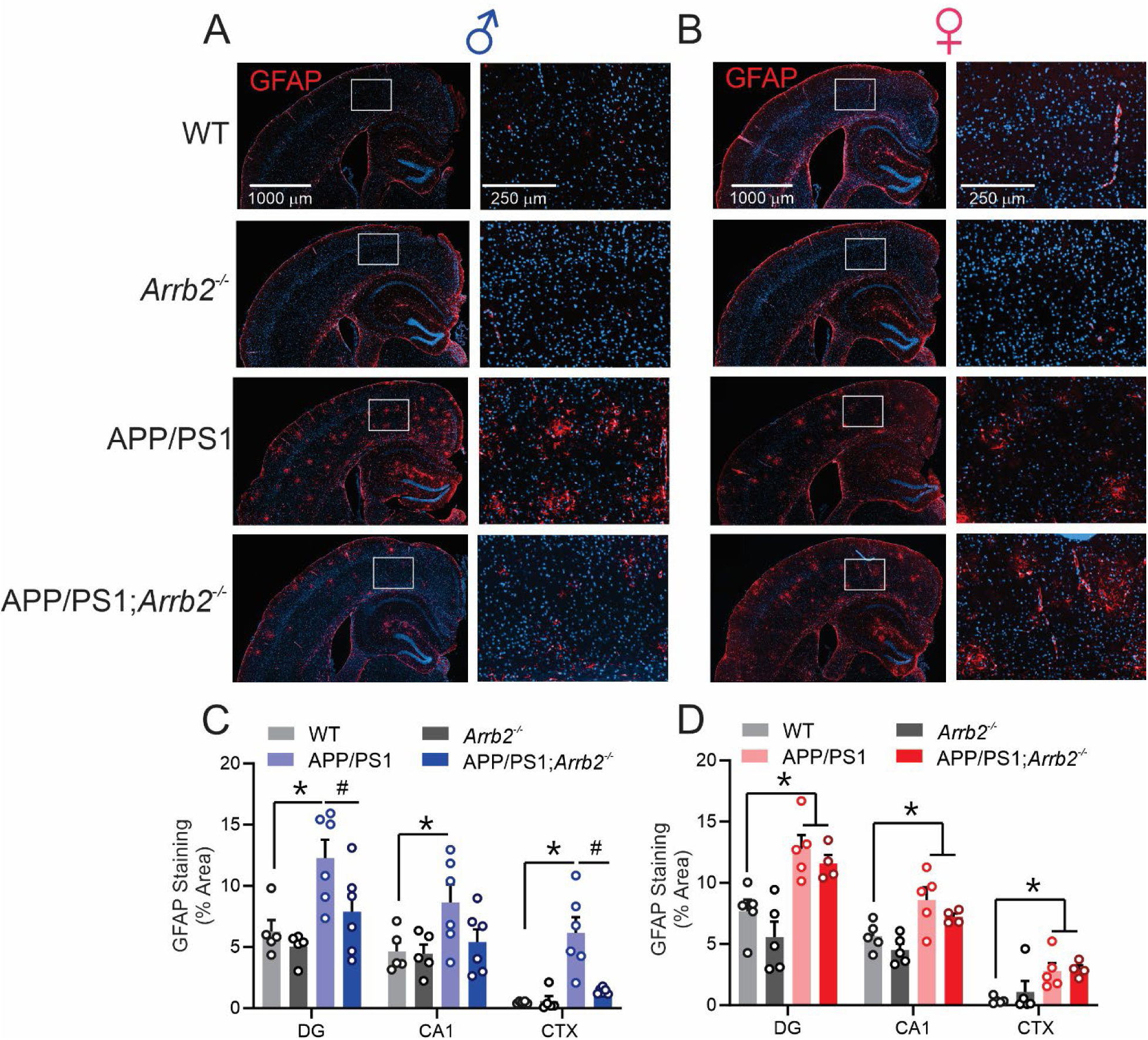
Effect of *Arrb2* deletion on astrogliosis in wild-type and APP/PS1 mice. **(A)** Representative images of Glial fibrillary acidic protein (GFAP) staining of prefrontal cortex and hippocampus from male WT, *Arrb2^-/-^,* APP/PS1and APP/PS1;*Arrb2^-/-^* mice. Corresponding boxed region is magnified in adjacent image. **(B)** Representative images of GFAP staining of prefrontal cortex and hippocampus from female WT, *Arrb2^-/-^,* APP/PS1and APP/PS1;*Arrb2^-/-^* mice. Corresponding boxed region is magnified in adjacent image. **(C)** Quantification of GFAP staining in 2 (hippocampus) or 4 (cortex) 450 x 450 µm regions of the dentate gyrus (DG), CA1 and cortex (CTX) of six brain slices per male APP/PS1 (*Arrb2^+/+^*) and APP/PS1;*Arrb2^-/-^* (*Arrb2^-/-^*) mouse. Data are presented as mean ± SEM (n = 5-6 mice per group). Statistical analysis was assessed using one-way ANOVA for each brain region for male and female mice individually followed by Tukey’s multiple comparisons test. *P < 0.05 versus region matched WT, # P< 0.05 versus region matched APP/PS1. **(D)** Quantification of GFAP staining in 2 (hippocampus) or 4 (cortex) 450 x 450 µm regions of the dentate gyrus (DG), CA1 and cortex (CTX) of six brain slices per female APP/PS1 (*Arrb2^+/+^*) and APP/PS1;*Arrb2^-/-^* (*Arrb2^-/-^*) mouse. Data are presented as mean ± SEM (n = 4-5 mice per group). Statistical analysis was assessed using one-way ANOVA for each brain region for male and female mice individually followed by Tukey’s multiple comparisons test. *P < 0.05 versus region matched WT, # P< 0.05 versus region matched APP/PS1.

### *Arrb2* deletion reduces astrogliosis and microgliosis in male but not female APP/PS1 mice

Astrogliosis and microgliosis are hallmark contributors to AD-associated neuroinflammation, with context-dependent effects on Aβ pathology^13^. GFAP immunoreactivity was elevated in cortex, dentate gyrus, and CA1 in both sexes of APP/PS1 mice relative to WT (Fig. 2A–D). *Arrb2* deletion normalized astrogliosis (glial fibrillary acidic protein positive, GFAP+) across all regions in males but had no effect in females (Fig. 2C, D). Ionized calcium binding adaptor molecule- 1 (Iba-1)+ cell counts were elevated in cortex and dentate gyrus of both sexes (Fig. 3A–D). *Arrb2* deletion reduced microgliosis in male cortex and dentate gyrus (Fig. 3C) but, conversely, increased Iba-1+ cell numbers in females (Fig. 3D). Thus, β-arrestin2 loss attenuates astrogliosis exclusively in male APP/PS1 mice and has opposing, sex-divergent effects on microgliosis in female APP/PS1 mice.

**Fig. 3:**
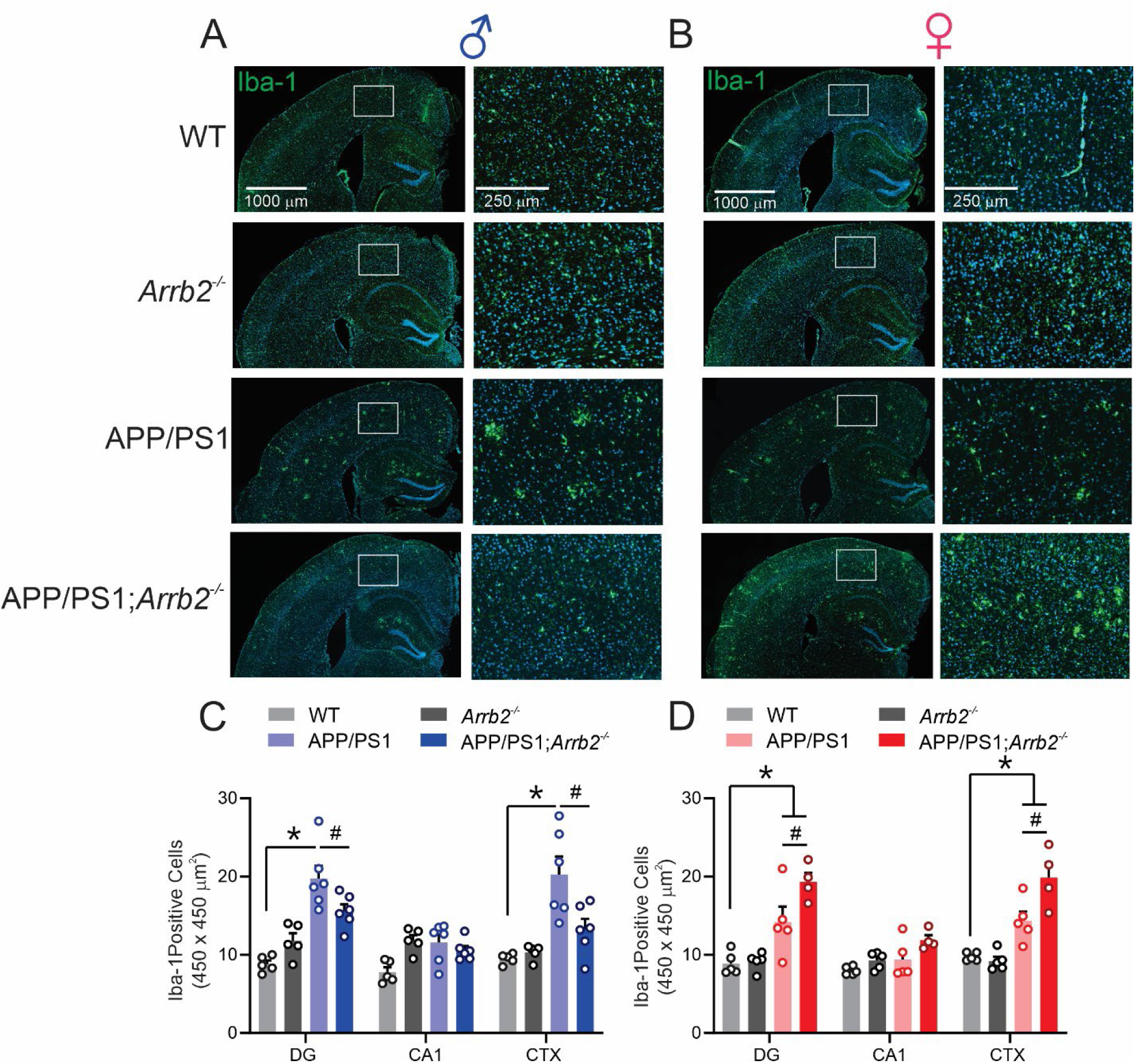
Effect of *Arrb2* deletion on microgliosis in wild-type and APP/PS1 mice. **(A)** Representative images of microglia-specific ionized calcium binding adaptor molecule 1 (Iba-1) staining in the prefrontal cortex and hippocampus from male WT, *Arrb2^-/-^,* APP/PS1and APP/PS1;*Arrb2^-/-^* mice. Corresponding boxed region is magnified in adjacent image. **(B)** Representative images of Iba-1 staining of prefrontal cortex and hippocampus from female WT, *Arrb2^-/-^,* APP/PS1 and APP/PS1;*Arrb2^-/-^*mice. Corresponding boxed region is magnified in adjacent image. **(C)** Quantification of Iba-1 cell counts in either 2 (hippocampus) or 4 (cortex) 450 x 450 µm regions of the dentate gyrus (DG), CA1 and cortex (CTX) of six brain slices per male WT, *Arrb2^-/-^*, APP/PS1 and APP/PS1;*Arrb2^-/-^* mouse. Data are presented as mean ± SEM (n = 5-6 mice per group). Statistical analysis was assessed using one-way ANOVA for each brain region for male and female mice individually followed by Tukey’s multiple comparisons test. *P < 0.05 versus region matched WT, # P< 0.05 versus APP/PS1. **(D)** Quantification of Iba-1 cell counts in 2 (hippocampus) or 4 (cortex) 450 x 450 µm regions of the dentate gyrus (DG), CA1 and cortex (CTX) of six brain slices per female WT, *Arrb2^-/-^*, APP/PS1 and APP/PS1;*Arrb2^-/-^* mouse. Data are presented as mean ± SEM (n = 4-5 mice per group). Statistical analysis was assessed using one-way ANOVA for each brain region for male and female mice individually followed by Tukey’s multiple comparisons test. *P < 0.05 versus region matched WT, # P< 0.05 versus APP/PS1.

### Effect of *Arrb2* deletion on cognitive behavior in WT and APP/PS1 mice

Given the sex-specific effects of *Arrb2* deletion on signaling, pathology, and gliosis, we assessed short-term spatial working memory (Y-maze), spatial learning (Morris water maze, MWM), cognitive flexibility (reversal MWM, RMWM), and recognition memory (novel object recognition, NOR) across genotypes and sexes. The Y-maze assesses short-term spatial working memory by exploiting a rodents’ innate tendency to avoid re-entering a recently visited arm; a successful alternation is scored when an animal sequentially enters all three arms (Fig. 4A). At 6 months, no genotype differences emerged in the Y-maze (Fig. 4B, C). At 9 months, *Arrb2* deletion reduced alternation frequency in WT mice of both sexes indicating that *Arrb2* deletion results in impaired spatial working memory in WT mice. In contrast, while zone alternations were also reduced in APP/PS1 mice of both sexes. *Arrb2* deletion rescued the alteration deficit in male but not female APP/PS1 mice (Fig. 4D, E).

**Fig. 4:**
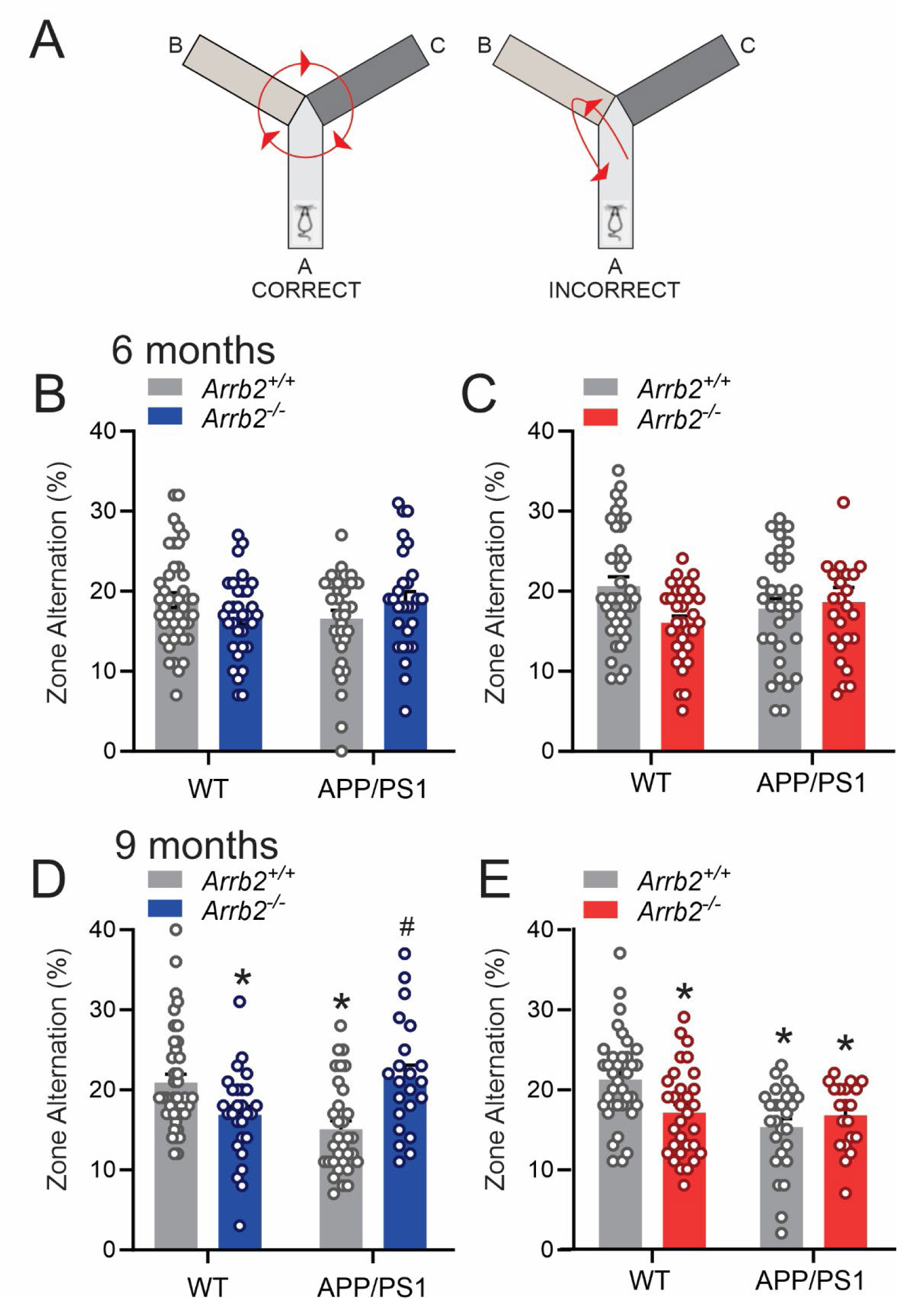
Effect of *Arrb2* deletion on working spatial memory in the Y-maze behaviour test for male and female WT and APP/PS1 mice at 6 and 9 month of age. **(A)** Schematic representation of correct and incorrect zone alterations in the open Y-maze behaviour test. A successful alteration is record when animals make consecutive entries into all three unique arms in overlapping triplet sets (e.g., B-C-A). **(B)** Correct zone alteration responses (%) for male WT, *Arrb2^-/-^*, APP/PS1, and APP/PS1;*Arrb2^-/-^* mice at 6 months of age. Data are presented as mean ± SEM (n = 25-41 mice per group). Statistical analysis was assessed using one-way ANOVA followed by Tukey’s multiple comparisons test. *P<0.05 versus sex matched WT. **(C)** Correct zone alteration responses (%) for female WT, *Arrb2^-/-^*, APP/PS1, and APP/PS1;*Arrb2^-/-^* mice at 6 months of age. Data are presented as mean ± SEM (n = 20-40 mice per group). Statistical analysis was assessed using one-way ANOVA followed by Tukey’s multiple comparisons test. *P<0.05 versus sex matched WT. **(D)** Correct zone alteration responses (%) for male WT, *Arrb2^-/-^*, APP/PS1, and APP/PS1;*Arrb2^-/-^* mice at 9 months of age. Data are presented as mean ± SEM (n = 22-40 mice per group). Statistical analysis was assessed using one-way ANOVA followed by Tukey’s multiple comparisons test. *P<0.05 versus sex matched WT. **(E)** Correct zone alteration responses (%) for female WT, *Arrb2^-/-^*, APP/PS1, and APP/PS1;*Arrb2^-/-^* mice at 9 months of age. Data are presented as mean ± SEM (n = 20-36 mice per group). Statistical analysis was assessed using one-way ANOVA followed by Tukey’s multiple comparisons test. *P<0.05 versus sex matched WT.

In the MWM/RMWM, both sexes of APP/PS1 mice showed prolonged escape latencies at 6 months, with *Arrb2*-null mice indistinguishable from WT (Fig. 5A-D). *Arrb2* deletion did not improve escape latency in APP/PS1 mice of either sex at this age (Fig. 5A–D). In the MWM at 6 months of age, both male and female *Arrb2^-/-^*, APP/PS1 and APP/PS1;*Arrb2*^-/-^ mice did not spend significantly different times in the target quadrant in the probe trial (Suppl. Fig. 1A and B). However, at 6 months of age male *Arrb2^-/-^*, APP/PS1 and APP/PS1;*Arrb2*^-/-^ mice spent less time in the target quadrant in the RMWM than WT controls, whereas there was no difference in target discrimination observed in female mice in the RMWM probe trial (Suppl. Fig. 1C and D). At 9 months, escape latency deficits persisted in APP/PS1 mice of both sexes, while *Arrb2*-null mice remained comparable to WT in this test (Fig. 5E–H). *Arrb2* deletion rescued MWM performance in male APP/PS1 mice only (Fig. 5E and F) and rescued RMWM performance in male but not female APP/PS1 mice as female PP/PS1;*Arrb2*^-/-^ mice failed to show significant improvement in the task from day 1 to day 3 (Fig. 5G and H). At 9 months of age all male and female APP/PS1 and APP/PS1 mice spent less time in the target quadrant during the MWM probe trail, whereas no target discrimination was observed in female mice in the RMWM probe trial, male APP/PS1 mice spent considerably less time the target quadrant (Suppl. Fig. 1G and H).

**Fig. 5:**
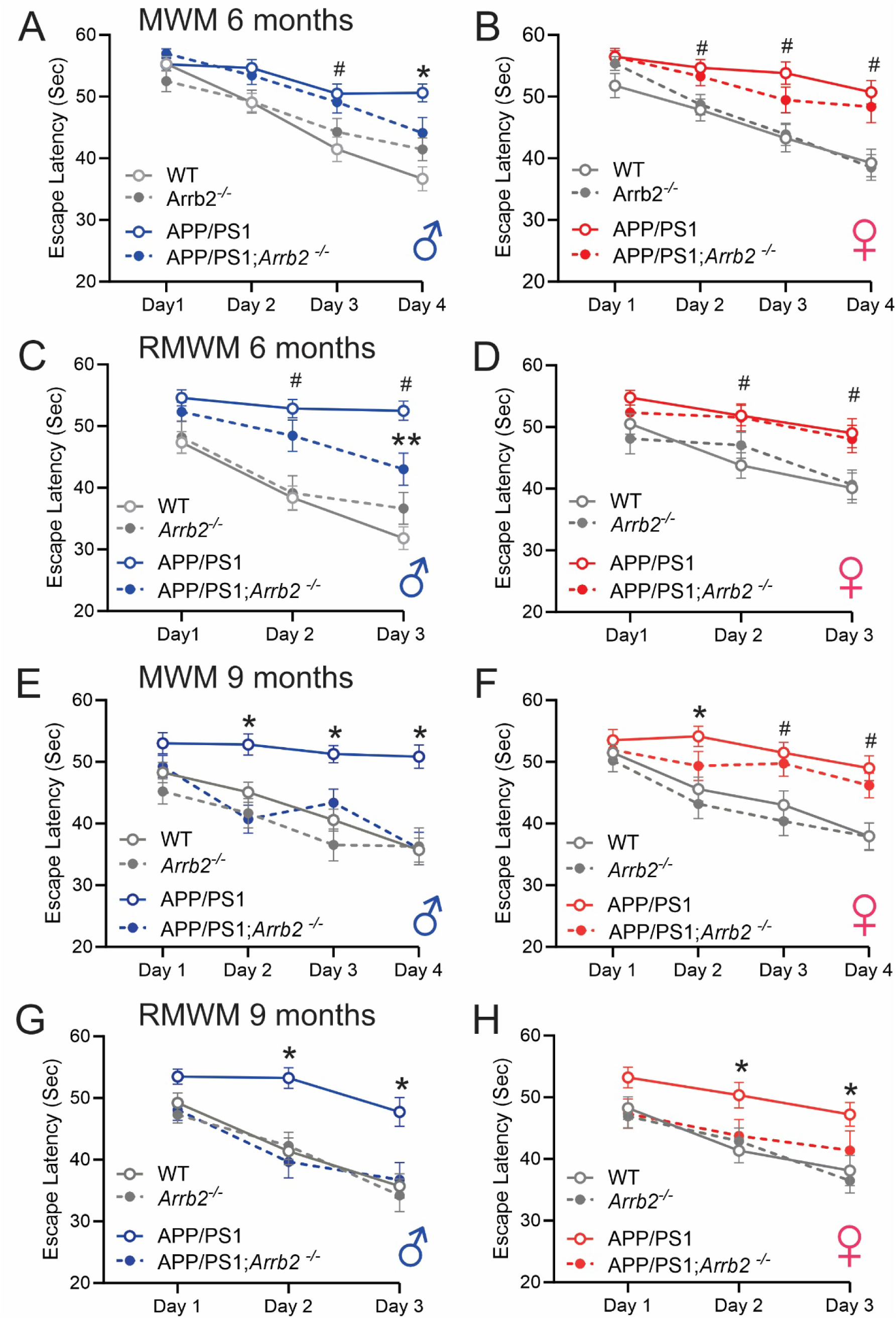
Effect of *Arrb2* deletion on male and female WT and APP/PS1 mice in the MWM and RMWM at 6 and 9 month of age. **(A)** Escape latency for male WT, *Arrb2^-/-^*, APP/PS1, and APP/PS1;*Arrb2^-/-^* mice in the MWM at 6 months of age. Data are presented as mean ± SEM (n = 25-39 mice per group). Statistical analysis was assessed using two-way ANOVA followed by Tukey’s multiple comparisons test (F(3,504)=16.75, P=0.0001). **(B)**Escape latency for female WT, *Arrb2^-/-^*, APP/PS1, and APP/PS1;*Arrb2^-/-^* mice in the MWM at 6 months of age. Data are presented as mean ± SEM (n = 21-36 mice per group). Statistical analysis was assessed using two-way ANOVA followed by Tukey’s multiple comparisons test. **(C)** Escape latency for male WT, *Arrb2^-/-^*, APP/PS1, and APP/PS1;*Arrb2^-/-^* mice in the RMWM at 6 months of age. Data are presented as mean ± SEM (n = 25-39 mice per group). Statistical analysis was assessed using two-way ANOVA followed by Tukey’s multiple comparisons test. **(D)** Escape latency for female WT, *Arrb2^-/-^*, APP/PS1, and APP/PS1;*Arrb2^-/-^* mice in the RMWM at 6 months of age. Data are presented as mean ± SEM (n = 21-36 mice per group). Statistical analysis was assessed using two-way ANOVA followed by Tukey’s multiple comparisons test. **(E)** Escape latency for male WT, *Arrb2^-/-^*, APP/PS1, and APP/PS1;*Arrb2^-/-^* mice in the MWM at 9 months of age. Data are presented as mean ± SEM (n = 22-39 mice per group). Statistical analysis was assessed using two-way ANOVA followed by Tukey’s multiple comparisons test. **(F)** Escape latency for female WT, *Arrb2^-/-^*, APP/PS1, and APP/PS1;*Arrb2^-/-^* mice in the MWM at 9 months of age. Data are presented as mean ± SEM (n = 20-36 mice per group). Statistical analysis was assessed using two-way ANOVA followed by Tukey’s multiple comparisons test. **(G)** Escape latency for male WT, *Arrb2^-/-^*, APP/PS1, and APP/PS1;*Arrb2^-/-^* mice in the RMWM at 9 months of age. Data are presented as mean ± SEM (n = 22-39 mice per group). Statistical analysis was assessed using two-way ANOVA followed by Tukey’s multiple comparisons test. **(H)** Escape latency for female WT, *Arrb2^-/-^*, APP/PS1, and APP/PS1;*Arrb2^-/-^* mice in the RMWM at 9 months of age. Data are presented as mean ± SEM (n = 20-36 mice per group). Statistical analysis was assessed using two-way ANOVA followed by Tukey’s multiple comparisons test. Tukey’s multiple comparisons test *P<0.05 APP/PS1 versus corresponding WT trial day. # P<0.05 APP/PS1 and APP/PS1;*Arrb2^-/-^* versus corresponding WT trial day. ** P<0.05 APP/PS1 versus corresponding APP/PS1;*Arrb2^-/-^* trail day.

In the NOR task, WT mice of both sexes discriminated novel from familiar objects at 6 months, while *Arrb2*-null and APP/PS1 mice of both sexes failed to do so (Fig. 6A and B). *Arrb2* deletion rescued discrimination in female, but not male, APP/PS1 mice at this age (Fig. 6A, B). By 9 months, *Arrb2* deletion rescued object discrimination in APP/PS1 mice of both sexes (Fig. 6C, D).

**Fig. 6:**
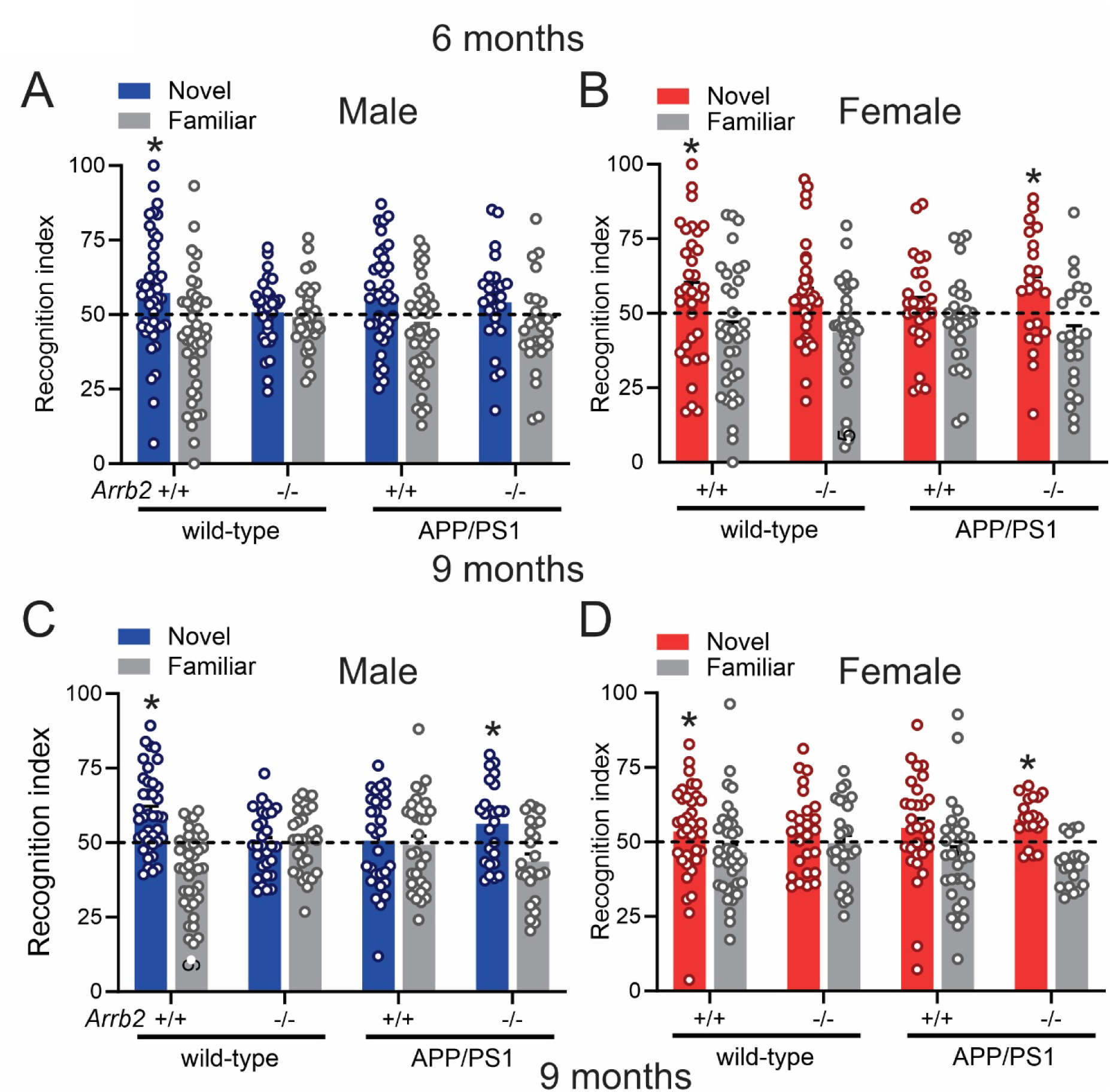
Effect of *Arrb2* deletion on recognition memory in male and female WT and APP/PS1 mice in the NOR test at 6 and 9 month of age. **(A)** The recognition index is shown for male WT, *Arrb2^-/-^*, APP/PS1, and APP/PS1;*Arrb2^-/-^* mice in the MWM at 6 months of age. Data are presented as mean ± SEM (n = 25-41 mice per group). Statistical analysis was assessed using two-way ANOVA followed by Newman-Keuls multiple comparisons post hoc test. *<0.05 versus familiar object. **(B)** The recognition index is shown for female WT, *Arrb2^-/-^*, APP/PS1, and APP/PS1;*Arrb2^-/-^* mice in the MWM at 6 months of age. Data are presented as mean ± SEM (n = 25-37 mice per group). Statistical analysis was assessed using two-way ANOVA followed by Newman-Keuls multiple comparisons post hoc test. *<0.05 versus familiar object. **(C)** The recognition index is shown for male WT, *Arrb2^-/-^*, APP/PS1, and APP/PS1;*Arrb2^-/-^* mice in the MWM at 9 months of age. Data are presented as mean ± SEM (n = 21-40 mice per group). Statistical analysis was assessed using two-way ANOVA followed by Newman-Keuls multiple comparisons post hoc test. *<0.05 versus familiar object. **(D)** The recognition index is shown for female WT, *Arrb2^-/-^*, APP/PS1, and APP/PS1;*Arrb2^-/-^* mice in the MWM at 9 months of age. Data are presented as mean ± SEM (n = 20-35 mice per group). Statistical analysis was assessed using two-way ANOVA followed by Newman-Keuls multiple comparisons post hoc test. * P<0.05 versus familiar object.

**Fig. 7.**
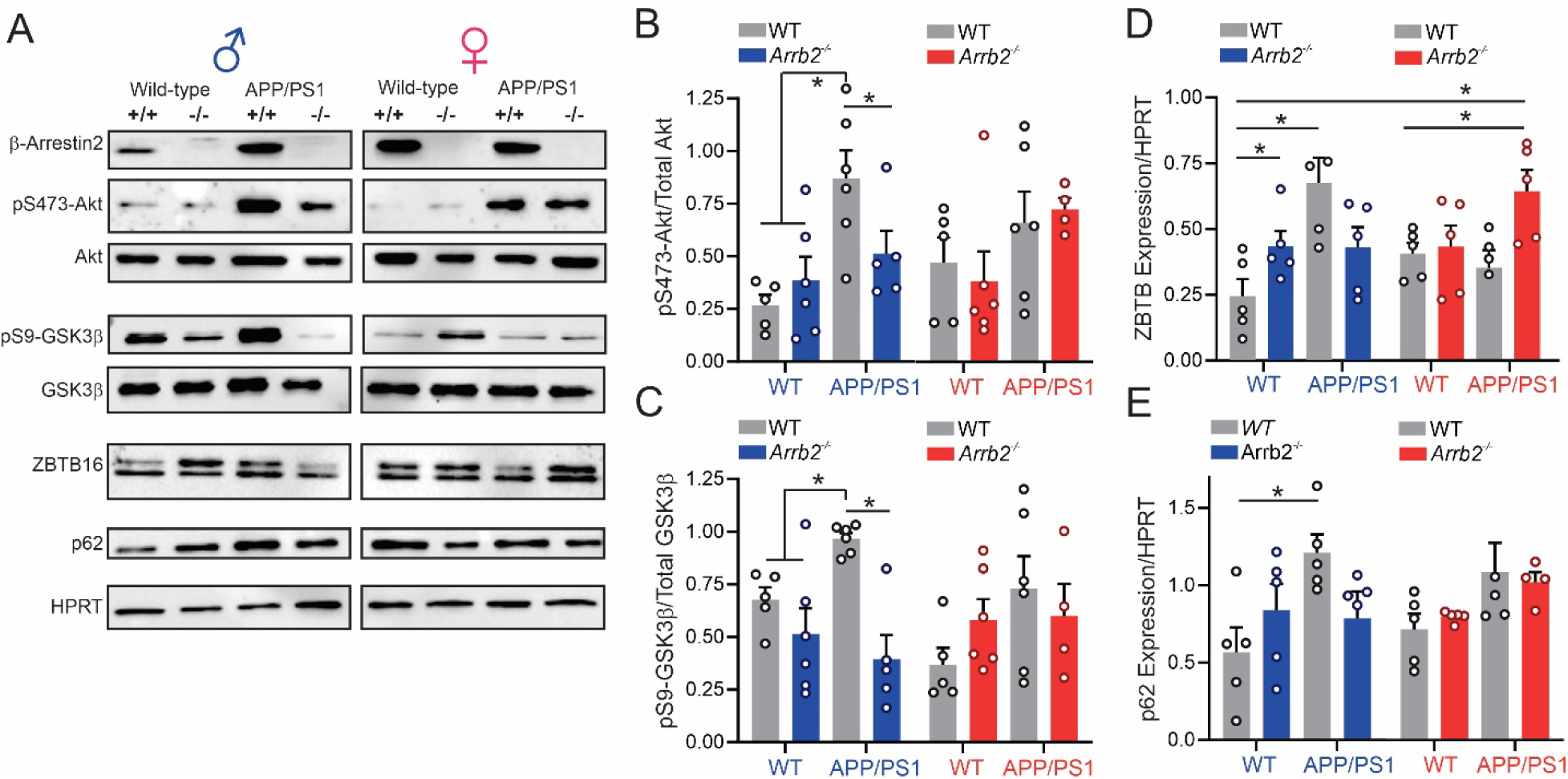
Effect of *Arrb2* deletion on pS473-Akt and pS9-GSK3β phosphorylation and ZBTB16 and p62 protein expression in male and female wild-type and APP/PS1 mice. **(A)** Representative immunoblots showing phosphorylation of Akt (pS473) and GSK3β (pS9) as well as protein expression of Akt, GSK3β, ZBTB16 and P62 in 9-month-old male and female wild-type (WT), *Arrb2^-/-^,* APP/PS1and APP/PS1;*Arrb2^-/-^*mice. N=4-6 independent experiments. (B) Quantitative analysis of pS473-Akt phosphorylation in cortical lysates from male and female WT, *Arrb2^-/-^,* APP/PS1and APP/PS1;*Arrb2^-/-^*mice normalized to Akt protein expression. Data are presented as mean ± SEM (n = 4-6 mice per group). Statistical analysis was assessed using two-way ANOVA followed by Tukey’s multiple comparisons test. *p<0.01 as indicated by the bars in the Figure. (C) Quantitative analysis of pS9-GSK3β phosphorylation in cortical lysates from male and female WT, *Arrb2^-/-^,* APP/PS1and APP/PS1;*Arrb2^-/-^* mice normalized to GSK3β protein expression. Data are presented as mean ± SEM (n = 4-5 mice per group). Statistical analysis was assessed using two-way ANOVA followed by Tukey’s multiple comparisons test. *p<0.05; ***p<0.01 as indicated by bars in the Figure. (D) Quantitative analysis of ZBTB16 phosphorylation in cortical lysates from male and female WT, *Arrb2^-/-^,* APP/PS1and APP/PS1;*Arrb2^-/-^* mice normalized to HPRT protein expression and male WT ZBTB16 protein phosphorylation. Data are presented as mean ± SEM (n = 4-5 mice per group). Statistical analysis was assessed using two-way ANOVA followed by Tukey’s multiple comparisons test. *p<0.05 as indicated by bars in the Figure (E) Quantitative analysis of p62 phosphorylation in cortical lysates from male and female WT, *Arrb2^-/-^,* APP/PS1and APP/PS1;*Arrb2^-/-^* mice normalized to HPRT protein expression and male WT p62 protein phosphorylation. Data are presented as mean ± SEM (n = 4-5 mice per group). Statistical analysis was assessed using two-way ANOVA followed by Tukey’s multiple comparisons test. **p<0.01, as indicated by bars in the Figure.

Together, these behavioral data indicate that while β-arrestin2 contributes to cognition in both sexes under normal conditions, *Arrb2* deletion predominantly rescues cognitive impairment in male APP/PS1 mice, with more limited, task-restricted rescue in females.

### β-Arrestin2 regulates male-specific Akt/GSK3β signaling and autophagy in APP/PS1 mice

Having found that Arrb2 deletion selectively reduces Aβ pathology, glial reactivity, and cognitive impairment in male APP/PS1 mice, we next asked what signaling mechanism might underlie this male-specific rescue. pS473-Akt and pS9-GSK3β phosphorylation are elevated in APP/PS1 and 5XFAD mice, and β-Arrestin2 is implicated in the regulation of Akt and GSK3β signaling^34,36,38–40,44^. We therefore asked whether these phosphorylation changes were sex-specific and β-arrestin2-dependent. In cortical lysates from 9-month-old mice, *Arrb2* deletion did not alter pS473-Akt or pS9-GSK3β in WT animals (Fig. 1A–C). However, both phospho-proteins were selectively elevated in male, but not female, APP/PS1 mice, and *Arrb2* deletion normalized these increases to WT levels (Fig. 1A-C). Because GSK3β regulates a ZBTB16-dependent autophagy pathway in APP/PS1, 3xTg-AD, and zQ175 HD mice^33–35,42,43^, we next assessed ZBTB16 and p62 expression. Both were selectively increased in male APP/PS1 cortex and restored to WT levels by *Arrb2* deletion (Fig. 1A, D, E), although ZBTB16 expression was independently elevated in *Arrb2*- /- mice alone. Together, these results indicate that β-arrestin2 drives male-specific Akt/GSK3β hyperactivation and autophagy suppression in APP/PS1 mice.

## Discussion

β-Arrestin2 signaling has been implicated in AD, PD, and FTD pathogenesis, but sex-specific differences in β-arrestin signaling had not previously been explored^27–30^, a notable gap given that AD disproportionately affects women and sex has been frequently neglected as a biological variable in preclinical research^4–6^. We find that both Aβ plaque density and soluble Aβ are reduced in the cortex of male, but not female, *Arrb2*-null APP/PS1 mice. We find that t pS473-Akt and pS9-GSK3β are selectively elevated in male APP/PS1 mice, and that *Arrb2* deletion reduces this phosphorylation and enhances GSK3β/ZBTB16-mediated autophagy in a male-specific manner. Consistent with previous studies, astrogliosis (GFAP) and microgliosis (Iba-1) are elevated in cortex and hippocampus of both sexes in APP/PS1 mice relative to wild-type and *Arrb2*^-/-^ controls^8,9,45^. However, *Arrb2* deletion reduces both GFAP and Iba-1 immunoreactivity in male APP/PS1 mice, while in females it fails to reduce astrogliosis and instead increases microglial numbers. This sex-dependent pattern extends to cognition: *Arrb2* deletion selectively rescues deficits in spatial working (Y maze) and learning (MWM) memories in males at 9 months but rescues recognition memory (NOR) performance in both sexes consistent with the broader role of β-arrestin2 in regulating signal transduction across a diverse array of GPCR subtypes. Together, these findings indicate that loss of β-arrestin2 preferentially normalizes signaling, pathology, glial reactivity, and cognition in male, rather than female, APP/PS1 mice.

Multiple GPCRs may converge on β-arrestin2-biased signaling to drive AD pathology. Here, *Arrb2* deletion produced male-specific attenuation of Aβ pathology was linked to reduced Akt/GSK3β phosphorylation and enhanced autophagy which phenocopies the effects of mGluR5 antagonism in APP/PS1 mice^33,34^. GPR3 and mGluR5 deletion reduces plaque deposition and improves memory in APP/PS1 mice, and a G protein-biased GPR3 knock-in selectively reduces Aβ pathology, indicating that G protein- and β-arrestin2-biased signaling are mechanistically separable therapeutic targets^9,10,46^. In P301S mice, β-arrestin2 loss reduces tau phosphorylation and rescues synaptic plasticity, implicating the protein in both Aβ and tau pathology^28^. *Arrb1* deletion likewise reduces Aβ accumulation and modestly improves MWM performance in APP/PS1 mice, consistent with both β-arrestin isoforms regulating γ-secretase via APH-1^30,32^.

Sex-specific GPCR/β-arrestin interactions are also documented beyond mGluR5. CRFR1 activation increases Aβ production through γ-secretase interaction^11^. CRFR1 overexpression produces AD-like pathology through divergent mechanisms by sex: Gαs-driven increases in tau and β-secretase1 phosphorylation and cognitive impairment in females versus β-arrestin2-biased signaling in males^11^. Stress-induced CRFR1/β-arrestin2 coupling is observed only in males^47^. However, this relationship is not strictly sex-invariant, as *Arrb1*-deficient males show elevated Aβ that is selectively reversible by PKA inhibition in females^47^. AT1AR knockout similarly reduces Aβ-induced cognitive impairment reflecting direct Aβ42 binding to astrocytic AT1AR that drives β-arrestin2-biased synaptotoxic signaling^8^. Finally, the G protein-biased β1AR agonist xamoterol reduces neuroinflammation and Aβ pathology in 5XFAD mice^12^. Together, this growing body of evidence positions β-arrestin2-biased signaling, as opposed to G protein-biased signaling, as a disease-promoting mechanism suggesting that G protein-biased signaling as a candidate target for safer, pathway-selective AD therapeutics.

Astrocytes and microglia critically shape both protective and pathological neuroglial signaling in AD^48–52^. Both cell types accumulate around Aβ plaques, and their reactivity can be modulated in a sex-dependent manner by GPCR activation or inhibition^8,9,48,53,54^, with reactive astrocytes capable of either exacerbating pathology via neurotoxic inflammatory signaling or conferring protection by suppressing it ^48,51,54,56^. We find that GFAP immunoreactivity is elevated in both male and female APP/PS1 mice, but that *Arrb2* deletion attenuates astrogliosis selectively in males, correlating with broader cognitive improvement. This male-specific effect parallels attenuated astrogliosis following mGluR5 antagonism or knockout in 5XFAD mice and astrocyte mGluR5-mediated Ca^2+^ dysregulation also promotes Aβ pathology and cognitive impairment in APP/PS1 mice suggesting that *Arrb2* deletion may act in part through loss of β-arrestin2-biased mGluR5 signaling^44,57^. A similar dependency is seen for AT1AR: astrocyte-specific AT1AR knockout reduces Aβ pathology and improves cognition in a β-arrestin2-dependent manner following Aβ42 oligomer exposure^8^. By contrast, a G protein-biased GPR3 mutant reduces Aβ pathology in APP^NL-G-F^ mice while paradoxically increasing both astrocytic and microglial hypertrophy^9^. This pattern echoes our own finding that, despite reducing microglial number in males, *Arrb2* deletion increases Iba-1+ microglia in female APP/PS1 mice. This raises the possibility that loss of β-arrestin2 signaling in females shifts glial GPCR signaling toward a G protein-biased, pro-hypertrophic state via Akt/GSK3β-independent mechanisms. Given the complexity of glial contributions to AD pathology, a more detailed dissection of β-arrestin2 function in astrocytes and microglia is warranted.

*Arrb2* deletion impaired Y-maze and NOR performance in both male and female wild-type mice, indicating that β-arrestin2 supports memory and learning under normal physiological conditions consistent with its requirement for mGluR1-dependent plasticity in CA3 and mGluR5-dependent plasticity in CA1 pyramidal neurons^37^. However, under AD-like pathophysiology, *Arrb2* deletion rescued cognitive impairment across the Y-maze, MWM, RMWM, and NOR in male APP/PS1 mice, likely reflecting reactivation of Akt/GSK3β-mediated autophagy and consequent Aβ clearance. In females, where *Arrb2* deletion did not alter Aβ pathology, cognitive rescue was restricted to NOR indicating that behavioral and pathological improvements can be dissociated to some extent. A similar dissociation was previously observed with the M1 muscarinic positive allosteric modulator VU0486846, which rescued memory in both sexes but reduced Aβ pathology only in females^54,58^. These findings underscore that β-arrestin2-dependent signaling in AD likely integrates across multiple GPCRs. For example, the expression of chemokine CXCL10 is increased in AD patients and AD mouse models and either antagonism or genetic deletion of its receptor, CXCR3, results in reduced concentrations of proinflammatory cytokines and attenuates the behavioral deficits in APP/PS1 mice^59^.

In conclusion, β-arrestin2 loss reduces Aβ pathology and broadly rescues cognition in male APP/PS1 mice, whereas in females it rescues a more limited subset of cognitive domains without altering Aβ pathology. These findings highlight the pleiotropic, sex-biased role of β-arrestin2 in GPCR signaling during AD progression and suggest that GPCR-targeted therapeutic strategies - whether agonist- or antagonist-based - should account for sex differences in G protein-versus β-arrestin2-biased signaling.

## Author contributions

Conceptualization: S.S.G.F. Methodology: F.P.A. Formal analysis: F.P.A., and S.S.G.F. Investigation: F.P.A., and T.L.C. and F.B. Writing: F.P.A. and S.S.G.F. Supervision and Project administration: S.S.G.F. Funding Acquisition: S.S.G.F.

## Data, code, and materials availability

All data and code needed to evaluate and reproduce the results in the paper are present in the paper. This study did not generate any new materials.

We would like to acknowledge the technical support provided by the Animal Behavior and Physiology Core and Louise Pelletier Histology Core facilities at the University of Ottawa.

## MATERIALS AND METHODS

### Reagents

Amyloid beta (Aggregated) Human ELISA Kit (KHB3491), goat anti-Rabbit (G-21234) and anti-mouse (G21040) IgG (H+L) Cross-Adsorbed HRP Secondary Antibody, rabbit anti-β-Amyloid (71-5800) and rabbit anti-β-Actin (PA1-183) were from Thermo Fisher Scientific. Rabbit anti- pSer^9^-GSK3β (9323) and mouse anti-GSK3β (9832) antibodies were from Cell Signaling Technology. Immunoprecipitation kit (206996), mouse anti-P62 (56416), rabbit anti-vinculin (129002) and rabbit anti-ZBTB16 (39354) were from Abcam. Rabbit anti-mGluR5 (AB5675) was from Millipore. β-Arrestin2-specific rabbit antibody was the gift of Dr. Jeffrey Benovic Jefferson University. Reagents used for western blotting were purchased from Bio-Rad and all other biochemical reagents were from Sigma-Aldrich.

### Animals

Breeding colonies were established from mice obtained from The Jackson Laboratory. We used B6;C3-Tg(APPswe,PSEN1dE9)85Dbo/Mmjax mice (strain 004462) mice. In addition, B6;C3-Tg(APPswe,PSEN1dE9)85Dbo/Mmjax (APP/PS1) mice were crossed with *Arrb2tm1Rjl*/J (strain 011130) to created heterozygous knockouts for *Arrb2* and either WT or hemizygous for APP/PS1. The heterozygous mice were bred to create our colony of homozygous knockout *Arrb2* with either WT or hemizygous APP/PS1 genotypes. Mice were group-housed, up to five animals per cage, with *ad libitum* access to food and water and were maintained under controlled environmental conditions on a 12-h light/12-h dark cycle at 24°C. Animals were moved to a reversed light cycle room at least 2 weeks prior to any behaviour study. All animal experimental protocols were approved by the University of Ottawa Institutional Animal Care Committee and were in accordance with the Canadian Council of Animal Care guidelines.

### Immunoblotting

Cortical tissue was lysed in ice-cold lysis buffer containing 50 mM Tris (pH 8.0), 150 mM NaCl, 1% Triton X-100, and 0.1% sodium dodecyl sulfate (SDS), supplemented with protease inhibitor cocktail and phosphatase inhibitors (10 mM NaF and 500 μM Na₃VO₄). Lysates were clarified by centrifugation at 15,000 rpm for 10 min at 4 °C. Protein samples were mixed with lysis buffer and 5× loading buffer containing β-mercaptoethanol, then heated at 90°C for 5 min. Samples were separated by SDS-PAGE and transferred onto nitrocellulose membranes (Bio-Rad). Membranes were blocked for 1 h at room temperature in Tris-buffered saline (pH 7.6) containing 0.05% Tween-20 (TBST) and 5% nonfat dry milk. Blots were then incubated overnight at 4 °C with primary antibodies diluted 1:1000 in TBST containing 2% nonfat dry milk. Following primary antibody incubation, membranes were incubated with appropriate HRP-conjugated secondary antibodies (anti-rabbit or anti-mouse) diluted 1:7000 in TBST containing 2% non fat dry milk for 1 h at room temperature. Protein bands were visualized and quantified using Clarity Western ECL substrate (Bio-Rad, 1705061).

### Determination of Aβ Oligomer by Sandwich ELISA

Levels of oligomeric Aβ were quantified using a sandwich ELISA approach as previously described^44^. Brains were dissected into right and left hemispheres, and one hemisphere was used for the measurement of oligomeric Aβ. Tissue samples were homogenized and centrifuged at 100,000 × g for 1 h at 4 °C to obtain soluble fractions enriched in Aβ oligomers. The resulting supernatant was collected and diluted 1:10 in the assay buffer provided with the kit. Quantification of oligomeric Aβ was performed using a commercially available sandwich ELISA kit (Thermo Scientific, #KHB3491) according to the manufacturer’s instructions. All samples were analyzed in triplicate. Protein concentrations were determined for each sample, and Aβ levels were normalized to total protein content.

### Immunostaining

One hemisphere of each brain sample was fixed in 4%-paraformaldehyde and then transferred to 70% ethanol for storage at 4°C. The samples were embedded in paraffin and then coronally sectioned through the hippocampus at a thickness of 5 µm. Sections were then incubated with the rabbit Aβ antibody at 1:200, Neuronal Nuclei (NeuN) antibody at 1:1500 for 30 minutes at room temperature and detected using an HRP conjugated compact polymer system. Slides were then stained using 3,3’-Diaminobenzidine (DAB) as the chromogen, counterstained with Hematoxylin, mounted and cover slipped. Slide were scanned using a Leica Aperio Slide scanner at 20× and the number of Aβ aggregates and NeuN positive cells were counted in representative 300 x 300 µm regions of interest (ROI) of cortex, as well as the dentate gyrus and CA1 (regions of the hippocampus. For immunofluorescence, paraffin embedded sections were deparaffinized and pre-treated using heat mediated antigen retrieval with EDTA buffer (pH 9.0). Slides were then rehydrated in TBST and blocked for 30 minutes with Rodent Block M (Biocare RBM9616). Sections were then incubated with Rabbit anti-GFAP (1:500) and Rabbit anti-Iba1 (1:1000) antibodies overnight at 4 ◦C. Sections were washed with TBST and then incubated with the Donkey anti-Rabbit Alexa Fluor ™ 568 and Goat anti-Mouse Alexa Fluor ™ 488 for 2 hours in the dark at room temperature. This was followed by incubation with a quencher (Vector TrueView Autofluorescence Quenching Kit SP-8400, Vector Labs) to decrease autofluorescence. Sections were then washed, incubated with 5 μg/ml of DAPI and coverslipped. Slides were scanned using a Leica Aperio Slide scanner (Wetzler, Germany) at 20× and the number of Iba1 positive cells and GFAP staining area (%) in immunohistochemistry sections were determined in representative 450*450 μm ROIs in the cortex, as well as the dentate gyrus and CA1 of the hippocampus. Experimenters were blinded to analysis and six sections per mouse were analyzed and for each section 2 ROIs in the hippocampus (CA1 and dentate gyrus) and 4 ROIs in the cortex were quantified using the cell counter tool in Image J^27,35^. This number of ROIs prevents the selection of only densely stained regions.

### Behavioral analysis

All animals were habituated in the testing room for a minimum of 30 minutes before testing. All behavioral tests were performed blindly and during the animal’s dark cycle.

### Spontaneous Y-maze

Spontaneous working memory was assessed using a Y-maze constructed from opaque acrylic consisting of three identical arms (30cm × 30 cm × 30cm) placed at 120° angles from each other, joined at a central point. The test relies on the animal’s natural curiosity and instinct to explore new environments rather than returning to familiar ones. Testing was conducted during the light phase under controlled illumination (60 lux). Mice were habituated to the testing room for 30 min before testing and were then allowed to freely explore the maze for 8 min. Behavioral activity was recorded using an overhead camera and analyzed using EthoVision 14 software to calculate spontaneous alternation performance, an index of working memory. A successful alteration occurs when the animal makes entries into all three unique arms in overlapping triplet sets. n alternation is defined as consecutive entries into all three unique arms (e.g., A → B → C, but not A → B → A). Alteration percentage is calculated as defined as: Percentage Alteration = (actual alterations/total entries – 2) x100.

### MWM and RMWM tests

The Morris water maze test was performed in a plastic pool (120 cm in diameter and 50 cm deep), filled with water made opaque by the addition of white nontoxic water-soluble paint, and maintained at 25 °C to prevent hypothermia. A clear escape platform (10 cm diameter) was placed 25 cm from the perimeter, submerged one cm beneath the surface of the water. Visual cues included a black “X” and square on the front and left walls of the maze room, respectively as spatial references. Mice were trained for 4 days (four trials per day and 15 mins between trails) to find the submerged platform at a fixed position from a random start point of the 4 equally spaced points around the pool. Each trial lasted either 60 seconds or until the mouse found the platform and mice remained on the platform for 15 seconds before being removed to their home cage. If the mice failed to find the platform within 60 seconds, they were guided to the platform by the experimenter. Escape latency was measured using Ethovision 10 automated video tracking software from Noldus and the average of four daily trials was plotted over the testing days. On day 5, the probe trial (a single trial of 60 seconds) was performed by removing the platform and allowing the mice to swim freely in the pool and recording the time spent in the target quadrant and the time spent in the outer perimeter of the pool (thigmotaxis time). RMWM task was initiated 24 hours after completion of MWM using the same paradigm as MWM, with 4 days’ acquisition and a probe trial. In the RMWM task, the platform was relocated to a new position and 3 days of testing was conducted as this was sufficient to detect group differences in learning in wild-type mice^27,28^.

### Novel object recognition test

Mice were placed in the empty box measuring 45 × 45 × 45 cm for 5 min and 5 min later, 2 identical objects were placed in the box 5 cm from the edge and 5 cm apart. Mice were returned to the box for 5 min and allowed to explore and the time spent exploring each object was recorded using a camera fed to a computer in a separate room and analyzed using Noldus Ethovision 10 software. Mice were considered to be exploring the objects if their snouts were within 1 cm of the object. Each experiment was repeated 24 hours after first exposure with one of the objects replaced with a novel object. Data was interpreted using recognition index which was as follows: time spent exploring the familiar object or the novel object over the total time spent exploring both objects multiplied by 100, and was used to measure recognition memory [TA or TB/ (TA + TB)]*100, where T represents time, A represents familiar object and B represents novel object^27,35^.

### Statistical analysis

Statistical analyses were performed using GraphPad Prism. Data are presented as mean ± SEM. Comparisons between groups for biochemical and immunoblotting experiments were conducted using one-way or two-way analysis of variance (ANOVA), as appropriate, followed by Tukey’s post hoc test for multiple comparisons.

## Abbreviations

Aβ: β-amyloid
AD: Alzheimer’s disease
AT1AR: angiotensin type 1A receptor
β2AR: β2-adrenergic receptor
CRFR1: corticotropin releasing factor receptor 1
GPCR: G protein-coupled receptor
GRK: G protein-coupled receptor kinase
GPR3: G protein receptor 3
Akt: protein kinase B
GSK3β: Glycogen synthase kinase-3β
mGluR5: metabotropic glutamate receptor 5
NAM: negative allosteric modulator
ZBTB16: Zinc Finger and BTB Domain Containing 16

---

**Supplemental Figure 1:**
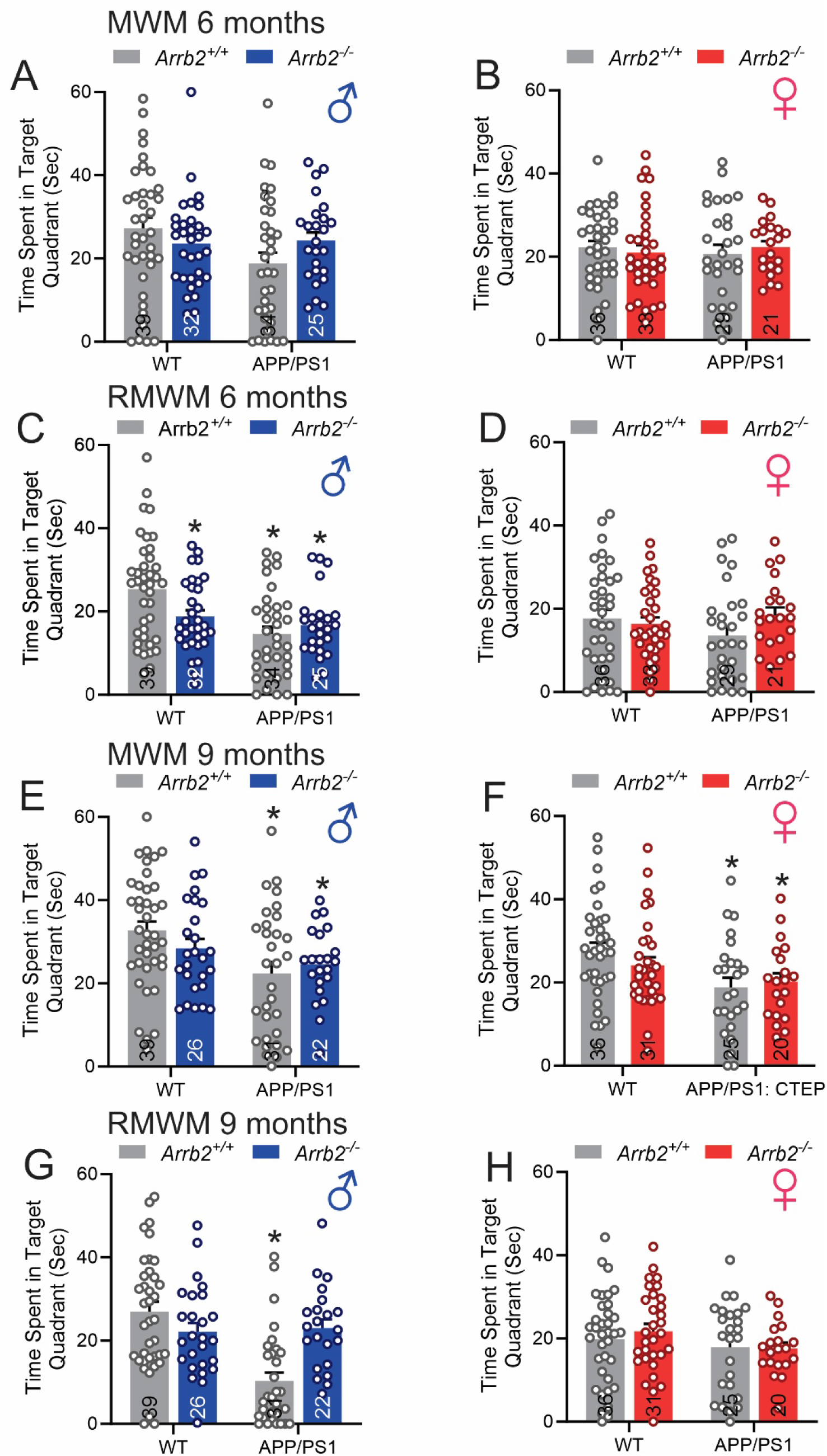
Effect of *Arrb2* deletion on time spent in the target quadrant male and female WT and APP/PS1 mice during MWM and RMWM probe trials at 6 and 9 month of age. **(A)** Time spent in the target quadrant for male WT, *Arrb2^-/-^*, APP/PS1, and APP/PS1; *Arrb2^-/-^* mice in the MWM at 6 months of age. Data are presented as mean ± SEM (n = 25-39 mice per group). Statistical analysis was assessed using one-way ANOVA followed by Tukey’s multiple comparisons test. **(B)** Time spent in the target quadrant for female WT, *Arrb2^-/-^*, APP/PS1, and APP/PS1;*Arrb2^-/-^* mice in the MWM at 6 months of age. Data are presented as mean ± SEM (n = 21-36 mice per group). Statistical analysis was assessed using one-way ANOVA followed by Tukey’s multiple comparisons test. **(C)** Time spent in the target quadrant for male WT, *Arrb2^-/-^*, APP/PS1, and APP/PS1;*Arrb2^-/-^* mice in the RMWM at 6 months of age. Data are presented as mean ± SEM (n = 25-39 mice per group). Statistical analysis was assessed using one-way ANOVA followed by Tukey’s multiple comparisons test. **(D)** Time spent in the target quadrant for female WT, *Arrb2^-/-^*, APP/PS1, and APP/PS1;*Arrb2^-/-^* mice in the RMWM at 6 months of age. Data are presented as mean ± SEM (n = 21-36 mice per group). Statistical analysis was assessed using one-way ANOVA followed by Tukey’s multiple comparisons test. **(E)** Time spent in the target quadrant for male WT, *Arrb2^-/-^*, APP/PS1, and APP/PS1;*Arrb2^-/-^* mice in the MWM at 9 months of age. Data are presented as mean ± SEM (n = 22-39 mice per group). Statistical analysis was assessed using one-way ANOVA followed by Tukey’s multiple comparisons test. **(F)** Time spent in the target quadrant for female WT, *Arrb2^-/-^*, APP/PS1, and APP/PS1;*Arrb2^-/-^* mice in the MWM at 9 months of age. Data are presented as mean ± SEM (n = 20-36 mice per group). Statistical analysis was assessed using one-way ANOVA followed by Tukey’s multiple comparisons test. **(G)** Time spent in the target quadrant for male WT, *Arrb2^-/-^*, APP/PS1, and APP/PS1;*Arrb2^-/-^* mice in the RMWM at 9 months of age. Data are presented as mean ± SEM (n = 22-39 mice per group). Statistical analysis was assessed using one-way ANOVA followed by Tukey’s multiple comparisons test. **(H)** Time spent in the target quadrant for female WT, *Arrb2^-/-^*, APP/PS1, and APP/PS1;*Arrb2^-/-^* mice in the RMWM at 9 months of age. Data are presented as mean ± SEM (n = 20-36 mice per group). Statistical analysis was assessed using one-way ANOVA followed by Tukey’s multiple comparisons test. *P<0.05 versus WT.

